# Integrating comparative modeling and accelerated simulations reveals conformational and energetic basis of actomyosin force generation

**DOI:** 10.1101/2022.09.22.508219

**Authors:** Wen Ma, Shengjun You, Michael Regnier, J. Andrew McCammon

## Abstract

Muscle contraction is performed by arrays of contractile proteins in the sarcomere. Serious heart diseases, such as cardiomyopathy, can often be results of mutations in myosin and actin. Direct characterization of how small changes in the myosin-actin complex impact its force production remains challenging. Molecular dynamics (MD) simulations, although capable of studying protein structurefunction relationships, are limited owing to the slow timescale of the myosin cycle as well as a lack of various intermediate structures for the actomyosin complex. Here, employing comparative modeling and enhanced sampling MD simulations, we show how the human cardiac myosin generates force during the mechanochemical cycle. Initial conformational ensembles for different myosin-actin states are learned from multiple structural templates with Rosetta. This enables us to efficiently sample the energy landscape of the system using Gaussian accelerated MD. Key myosin loop residues, whose substitutions are related to cardiomyopathy, are identified to form stable or metastable interactions with the actin surface. We find that the actin-binding cleft closure is allosterically coupled to the myosin core transitions and ATP-hydrolysis product release from the active site. Furthermore, a gate between switch I and switch II is suggested to control phosphate release at the pre-powerstroke state. Our approach demonstrates the ability to link sequence and structural information to motor functions.

**Significance Statement:** Interactions between myosin and actin are essential in producing various cellular forces. Targeting cardiac myosin, several small molecules have been developed to treat cardiomyopathy. A clear mechanistic picture for the allosteric control in the actomyosin complex can potentially facilitate drug design by uncovering functionally important intermediate states. Here, integrating Rosetta comparative modeling and accelerated molecular dynamics, we reveal how ATP-hydrolysis product release correlates with powerstroke and myosin tight binding to actin. The predicted metastable states and corresponding energetics complement available experimental data and provide insights into the timing of elementary mechanochemical events. Our method establishes a framework to characterize at an atomistic level how a molecular motor translocates along a filament.

---

**M**any physiological processes are driven by mechanical force generated through myosin-actin interactions, such as muscle contraction, vesicle trafficking, and membrane deformation (1, 2). It is remarkable that actomyosin carries out these diverse functions through a conserved mechanochemical cycle, in which the chemical energy from ATP hydrolysis is used to generate force via myosin motor domain conformational changes (3, 4). The physical properties of the cycle are tuned to accomplish a variety of cellular tasks for different myosin isoforms (5, 6).

In muscle, the basic contractile apparatus is formed primarily by myosin (thick) and actin (thin) filaments, which slide past each other to contract the muscle fiber (7). The interaction between myosin head and actin powered by ATP results in the cross-bridge formation (8). Hypertrophic cardiomyopathy (HCM) and dilated cardiomyopathy (DCM), two leading causes of cardiac death, can often be due to mutations in sarcomeric proteins (9). HCM mutations cause hypercontractile state of the sarcomere whereas DCM mutations are linked to ventricular dilatation and loss of systolic function. Studies have shown that HCM and DCM mutations in myosin give rise to changes in the basic physical parameters of the mechanochemical cycle (10, 11) or the number of available myosin heads to interact with actin (12, 13). Several small molecules targeting the myosin motor domain (14–16) have been developed to treat these cardiac diseases, but clear molecular bases for the drug effects remain to be established (17).

The actomyosin functional cycle includes two major stages: a force-generating stage in which myosin swings its lever arm and stays engaged with actin (Fig. 1A), and a recovery stage in which myosin returns the lever arm to a primed configuration and stays detached from the actin filament. The force-generation stage involves a few key processes, i.e., formation of a tight myosin-actin interface, actin-binding cleft closure, lever arm swing, and ATP hydrolysis product (phosphate and ADP) release. Despite extensive structural, biochemical, and single-molecule studies (see reviews (6, 18)), the causality and ordering of these events are difficult to characterize. Recent cryo-EM structures provide atomistic views of different myosin-actin isoforms at the strongly bound rigor state (19–22), in which no nucleotide is bound to the myosin active site. Fewer mechanistic details are known for the early binding events and the powerstroke transition. So far no actomyosin structures have been solved for the pre-powerstroke (PPS) state due to its weak binding nature. Although it is generally believed that myosin attachment to actin leads to phosphate (Pi) release and powerstroke, the timing of Pi release and lever arm motion is still equivocal. A FRET study on myosin V (23) suggested that the initial contact triggers a fast lever arm motion, followed by a slower stroke swing after Pi release and before ADP release. Later on, a high-resolution optical tweezers experiment on cardiac myosin (24) demonstrated that the powerstroke rate is much faster than the estimated Pi release rate.

**Fig. 1.**
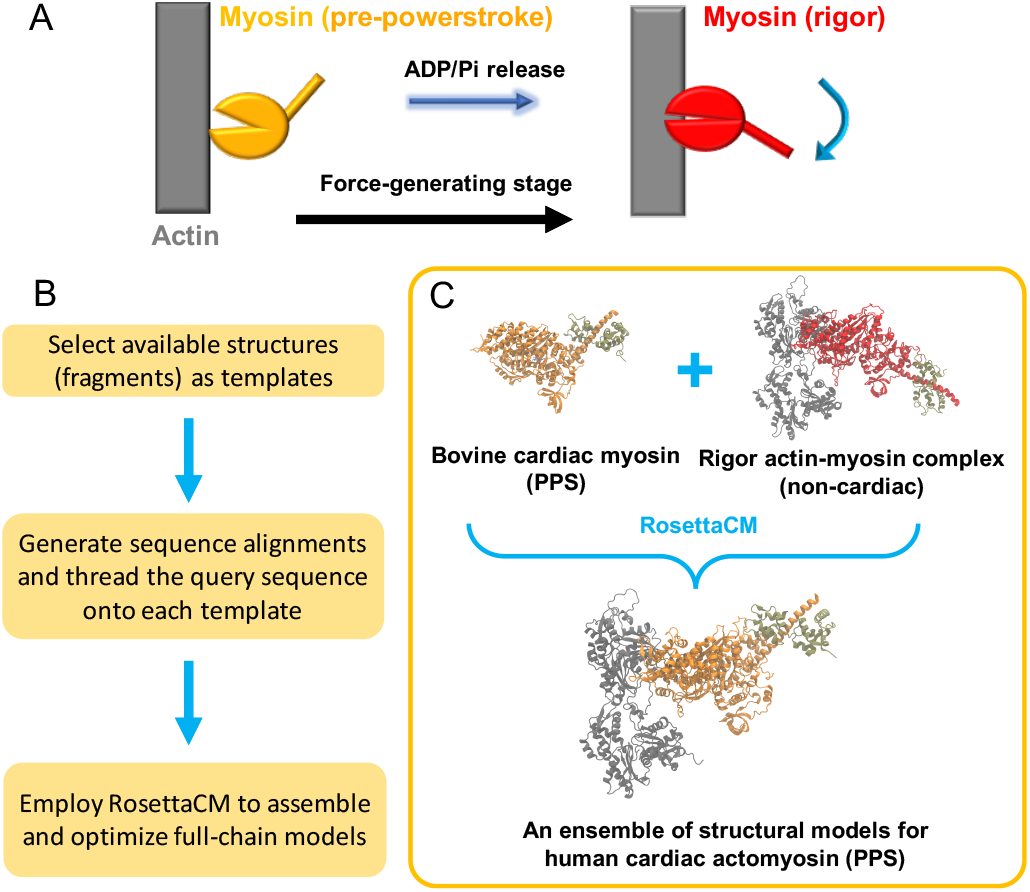
Modeling of multiple actomyosin states. A. Simplified scheme for the forcegeneration stage, during which the myosin is transitioned from the pre-powerstroke (orange) to the rigor (red) states. B. The flowchart of comparative modeling with Rosetta. C. A specific case in which structural models for actin-bound human cardiac myosin at the PPS state are obtained by using two templates – 5N69 (bovine cardiac muscle) and 5H53 (rabbit skeletal muscle). The former is a crystal structure of an isolated myosin at the PPS state (orange) whereas the latter is a cryo-EM structure of the actin (gray) – myosin (red) complex at the rigor state. ADP and Pi are explicitly incorporated in the active site of the PPS actomyosin models.

A clear atomistic-level picture for the allosteric regulation encoded in the motor domain is crucial to understanding how small-molecule drugs impact myosin function, and to help design and optimize molecules targeting allosteric sites based on key intermediate states. Recently a machine learning based method AlphaFold2 has successfully demonstrated the ability in predicting protein folds (25) and multimeric interfaces (26) given a query sequence. However, this approach is limited in studying the actomyosin cycle, due to its inability to handle ligand and mutation effects, as well as multiple conformational states and transitions (27, 28). Developed over decades, all-atom molecular dynamics (MD) simulations have become powerful in studying the mechanisms of biomolecular machines (29–31). By combining Rosetta comparative modeling and Gaussian accelerated molecular dynamics (GaMD) (32), we develop a computational approach that characterizes the conformational ensembles of actomyosin at different ligand states. Our results reveal the coupling between actin-binding cleft closure and structural rearrangement at the active site. The population distribution of the PPS state along myosin conformational coordinates is shifted to that of the rigor state, as a consequence of hydrolysis product release. The predicted actin-myosin interactions agree with the existing rigor cryo-EM structures and mutagenesis studies. Our work highlights the principles underlying force-producing mechanisms of myosin-actin systems.

## Results

### Ensemble structures of the myosin-actin complex at different states learned by comparative modeling

Due to the lack of high-resolution intermediate structures in the cardiac crossbridge cycle, we first applied comparative modeling to probe the conformational space of the myosin-actin complex. The approach combines structural information from multiple templates and uses RosettaCM (33) to build the complex (Methods). The effects of ligands (e.g. ADP and Pi) are explicitly considered. In Fig. 1B and 1C, the case to build the actin-bound pre-powerstroke (PPS) state models is illustrated, as we combined the information from two templates (an isolated PPS myosin structure and a rigor actin-myosin complex). Since no experimental structure is available for the PPS actomyosin, our modeling enables us to study the effects of hydrolysis product release on myosin-actin interactions. We also built the rigor and ADP-bound myosin-actin complex structures based on different templates.

The models were ranked by the Rosetta score function and the top conformations were selected for each state and then clustered. The most populated conformations for PPS and rigor states are shown in Fig. 2A and 2B. The lever arm at the rigor state exhibits a big downward swing compared with that at the PPS state. The two ensembles both exhibit structural variations at the myosin-actin interface. The myosin loop conformations are overlaid in Fig. 2C and 2D after aligning the models to the upper actin molecule. Among the key myosin loops, loop 2 displays large fluctuations, and loop 4 exhibits large positional drifts. The cardiomyopathy (CM) loop, which forms hydrophobic contacts with the upper actin, has relatively smaller variations. The average contact area between the CM loop and actin is 351 Å^2^ for the rigor models, larger than the 336 Å^2^ average area for the PPS models. This is consistent with the fact that PPS is a weaker actin-binding state. The helix-loop-helix (HLH) motif of the L50 subdomain also forms important contacts with actin. The average contact area between the HLH motif and actin is 516 Å^2^ for the rigor models, smaller than the number (581 Å^2^) for the PPS models. We note that the above numbers based on comparative modeling do not represent the physical ensemble averages. To validate these structures, we carried out enhanced sampling simulations starting from the Rosetta models of each state. Previously short conventional MD was shown to improve Rosetta sampling in an iterative refinement protocol guided by cryo-EM density (34).

**Fig. 2.**
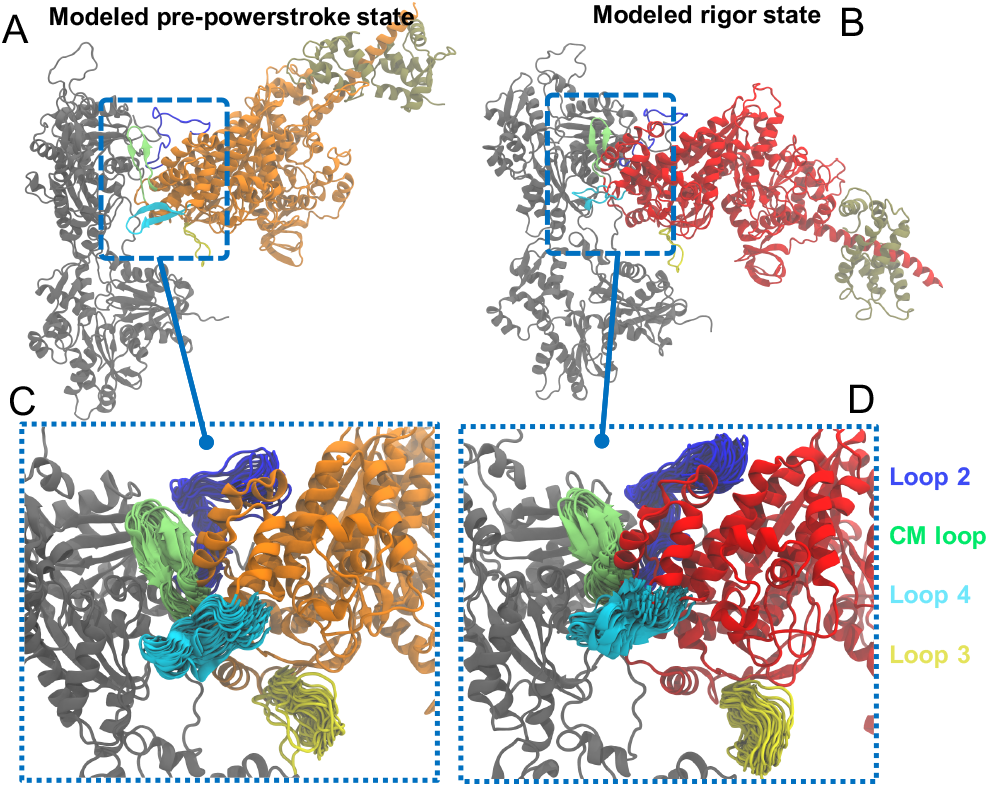
Structural ensembles at two different myosin states resulted from Rosetta comparative modeling. A and B: centroid structures for PPS and rigor actin-myosin complex, respectively. A centroid structure was obtained by clustering all the Rosetta models for a given state. C and D: a closer look at the key myosin loop regions with all the models overlaid at each state. To make the representation clear, the rest of the complex only displays the centroid structure. Actin is shown in gray whereas myosin is colored in orange (PPS) or red (rigor).

### Enhanced sampling MD simulations reveal important myosin-actin interactions

The conformational states from Rosetta do not reflect thermodynamic equilibrium and some of the states might locate in the high energy regions of the free energy landscape. Next, energy minimization and MD equilibration were performed for each initial Rosetta model. Multiple GaMD simulations were continued from each of the successful initial equilibration simulations at different states (pre-powerstroke, ADP-bound, and rigor, see Methods). By combining and reweighting the trajectories to recover the equilibrium distributions, 1D or 2D free energy profiles were projected along representative coordinates. Our results illustrate major features of the interface at different states in the following.

We first inspected the interactions between myosin and actin at the rigor state. The key myosin motifs identified above exhibit dynamic features while remaining attached to actin (Fig. 3 and Movie 1). As shown in cryo-EM structures (20–22), the myosin CM loop forms stable hydrophobic interactions with the upper actin surface (Fig. 3A), e.g., as evidenced by the distance trace (Fig. 3D) between V406 (CM loop) and A26 (actin) and the probability distribution (Fig. S1A). The related mutations (V406M, A26V) have been found to cause HCM (35). E371 on Loop 4 forms a metastable electrostatic interaction with K328 on actin as shown in Fig. 3B. Here the most probable distance between CD atom of E371 and NZ atom of K328 is 3.8 Å (Fig. 3B and S1B). This is consistent with a recent cryo-EM structure of cardiac rigor actomyosin in which a very similar loop 4 – actin contact was established (22). Loop 2, which is not visible in previous structural studies, is indeed flexible in our simulations. K635 is found to form electrostatic interactions with the negatively-charged N-terminus of the upper actin, e.g. E4 (Fig. 3C). Variation of this residue is linked to DCM phenotype (36). The interaction between K635 and E4 of actin is rather transient with the most populated distance at 3.8 Å and the second most populated state at around 6.9 Å (Fig. 3F and S1C), indicating that this interaction contributes less to strong actin-binding states.

**Fig. 3.**
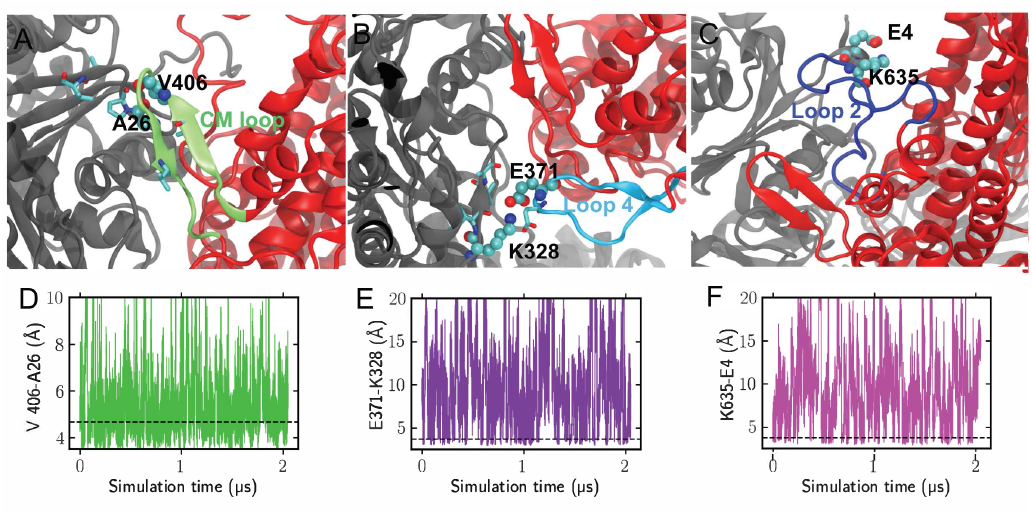
GaMD simulations reveal key myosin loop interactions with actin at the rigor state. A-C. Representative conformations for the CM loop, loop 4, and loop 2, respectively. D-F. Distances between the highlighted residues as a function of the accumulated simulation time. The dashed horizontal black lines represent the most probable distances, which are also demonstrated in the probability distributions shown in Fig. S1.

We evaluated the simulation results by comparing them to the experimental data. A representative rigor interface conformation (from a top cluster) was fit to a recently published rigor actomyosin cryo-EM density (EMD-22335, 3.8 Å resolution) (22). The CM loop, Loop 4, and HLH motif, along with the nearby actin surface, fit nicely into the density as shown in Fig. S2. Moreover, the MD structures were aligned to published myosin structures (21, 22, 37) and the motor core RMSD from different PDBs are listed in Table S1. The rigor simulations have an average RMSD of 2.3 Å from rigor PDBs (7JH7 and 6X5Z), which is relatively small compared with the 3.6 Å from a PPS PDB (37) (5N69). On the other hand, the average RMSD of the PPS simulations is 3.5 Å from the rigor PDBs and 1.5 Å from the PPS PDB. More details about the PPS actomyosin interface are provided in SI.

### Dynamic coupling between actin binding and myosin powerstroke

To probe if there is communication between remote sites across myosin, we measured the actin-binding cleft width and the myosin HF helix rotation (Fig. 4A) for our simulations at different states. The cleft width determines the separation between the upper and lower 50K domains. The relative rotation of the HF helix to *β*5 of the transducer, which is correlated with the transducer twisting (Fig. 4B), is a central process in myosin force generation. The structural ensemble of the PPS simulations is compared with that of the rigor simulations, as illustrated by the 2D free energy profiles projected along the cleft width and the HF rotation coordinates (Fig. 4D and 4E). For the rigor state, the rotation angle (x-axis) fluctuates near the basin around 5 ∼ 10 degrees, and is smaller than the one at the PPS state, which fluctuates around 20 ∼ 25 degrees. This rotation of the HF helix from PPS to the rigor state facilitates the N-terminal subdomain motion relative to the U50 subdomain (Fig. 4C). In the meantime, the cleft width decreases from 20 Å to 15 Å by comparing the y-axis values at the energy minima in the two plots. We thus suggest that the cleft closure rearranges the myosin core helices in the U50 and L50 subdomains, allowing transducer twisting to occur.

To illustrate the major changes in the actin-myosin interactions before and after the powerstroke, we calculated the contact areas between key motifs and actin and evaluated the contributions of individual motifs. 2D free energy profiles were obtained using contact areas of the CM loop-actin and HLH-actin interfaces. At the rigor state (Fig. S3A) the most probable contact areas of the two interfaces are 390 Å^2^ and 609 Å^2^, respectively, showing that the L50 subdomain contributes a larger contact area than the CM loop. At the PPS state (Fig. S3B) the most probable contact areas for the CM loop-actin and HLH-actin interfaces are 377 Å^2^ and 688 Å^2^, respectively. Interestingly, a few of metastable states (labeled by red circles in Fig. S3B) were located on the PPS energy landscape. These states display a much smaller contact area (*<*200 Å^2^) between the CM loop and actin, whereas the contact between the HLH motif and actin remains strong (*>*500 Å^2^). Overall, the CM-actin interface is weaker at the PPS state and the HLH motif stays attached to actin at both states. This observation indicates that the L50 subdomain forms stable interactions with actin first during initial binding, followed by the engagement between the CM loop and actin, which is coupled to the relative motion between the lower and upper domains (38).

**Fig. 4.**
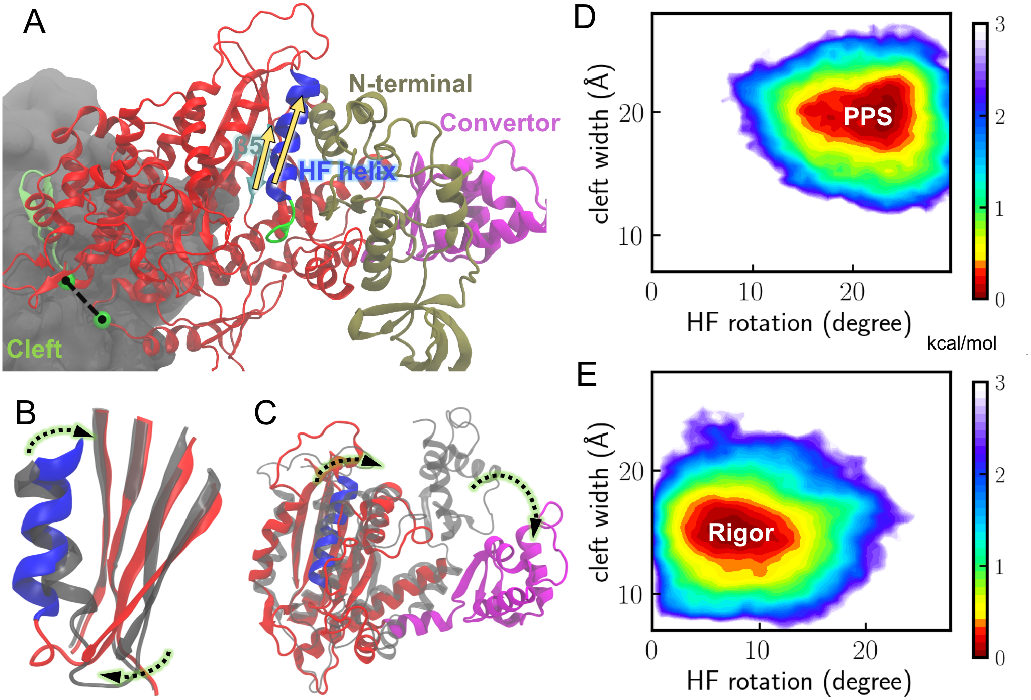
Correlations between actin-binding cleft closure and conformational changes near the ATP binding site. A. A representative structure of the most populated cluster at the rigor state. The cleft width is measured here by the distance between V417 (precedes the CM loop) and K542 (in the helix-loop-helix motif). The C*α* atoms of the two residues are shown in green spheres. The relative rotation between *β*5 of the transducer and the HF helix of the N-terminus subdomain is measured by the crossing angle between the two corresponding vectors (yellow arrows). B. HF helix rotation and transducer twisting from the PPS (colored in gray) to the rigor state (with HF colored in blue and transducer in red). C. Rotation of the converter (colored in magenta) during the powerstroke. The PPS myosin is colored gray. D and E. 2D free energy profiles for the PPS and rigor states are calculated along two coordinates (HF rotation angle and cleft width). The free energy profiles reflect the equilibrium distributions of the system (details in SI Methods).

### Free energy profiles reveal active site rearrangements induced by hydrolysis product release

Following actin binding, the interface motion and binding cleft closure promote the transitions of motor core helices, which enable transducer twisting (Fig. 4). We reasoned that the different ATP binding states (ADP+Pi, ADP, and empty) might shift the populations of the motor domain conformational states via allosteric regulations. By comparing the binding site dynamics at different states and estimating the free energy profiles along two distance coordinates, the population shifts upon product release are shown in Fig. 5.

**Fig. 5.**
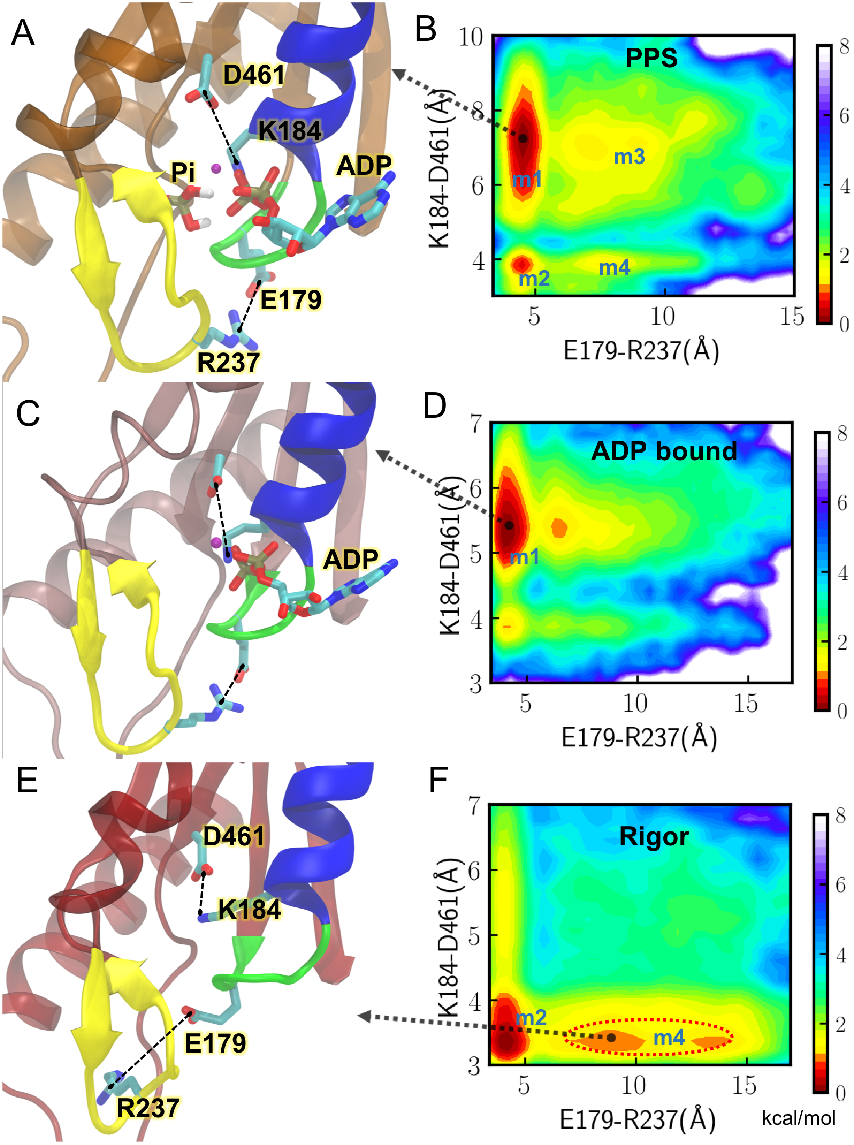
Pi and ADP release change the coordination of key motifs at the binding site. Representative conformations for the binding site at the PPS (ADP+Pi bound), ADP-bound, and rigor states are shown in panels A, C, and E, respectively (P loop: green, switch I: yellow, HF helix: blue). Two measured distances (E179-R237 and K184-D461) are highlighted by dotted black lines. B, D, and F. Free energy profiles along two distance coordinates at the PPS, ADP-bound, and rigor states, respectively. A black arrow points to a representative conformation at a given energy basin on the landscape. m1, m2, m3, and m4 are used to label local energy basins.

The feature changes upon Pi release are demonstrated in Fig. 5A-D. At the PPS state, four local minima m1 – m4 are featured in the free energy landscape generated using the P loop – switch I and P loop – *β*5 distances (Fig. 5A and 5B). The lowest energy state is m1, in which E179 (P loop) forms an electrostatic interaction with R237 (switch I). In the meantime, K184 (P loop) forms interactions with *β* and *γ* phosphates and is disengaged from D461, which locates at the end of *β*5 of the transducer. At the ADP-bound state, the distance between K184 and D461 at the lowest energy state m1 decreases to 5.5 Å (Fig. 5D) from the 7 Å at PPS (Fig. 5B) suggesting that Pi release is linked to the transducer transition through this interaction. This movement also affects the switch II motion, since D461 is in the loop connecting *β*5 to switch II.

A tunnel is found to open between switch I and switch II at the PPS state, as indicated by the opening of the salt bridge between R243 (switch I) and E466 (switch II) shown in Fig. 6C and Movie 2. The distance between the two residues at the PPS state renders a bimodal distribution (Fig. 6B), which favors the closed state (Fig. 6A) over the open tunnel (Fig. 6C). The P loop – switch I interactions remain stable during Pi release as suggested by the small 4 Å distance between E179 and R237 at m1 of both PPS and ADP-bound states. Thus our results favor a Pi release mechanism via the “back door” (39), which is achieved by the switch II movement. The “side door” release mechanism (38) is less likely to happen because of the strong interactions between the P loop and switch I in m1 at both PPS and ADP-bound states.

**Fig. 6.**
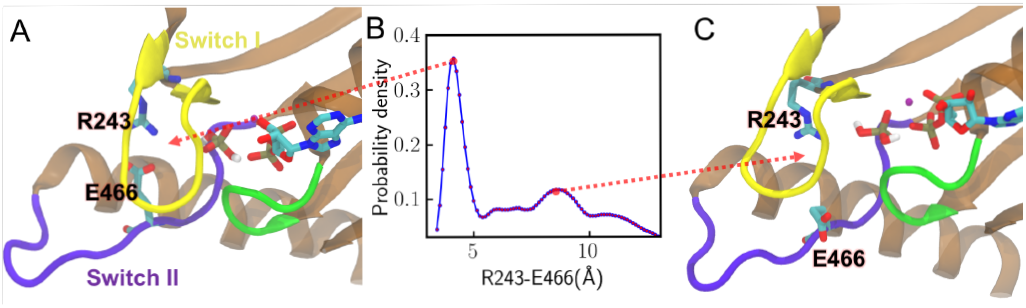
A gate for Pi release formed between switches I and II at the PPS state. A. The most probable conformation of switch I and II. B. Probability distribution for the distance between R243 (switch I) and E466 (switch II). C. The representative conformation at the second peak in the distribution. The red dotted lines point to the respective structures at the two peak positions.

By comparing the results for ADP-bound and rigor actomyosin, the effect of ADP release on the motor is demonstrated in Fig. 5C-F. The rigor conformational distribution is very different from those at the PPS and ADP-bound states. At the ADP-bound state, K184 on the P loop coordinates with *β*-phosphate, while E179 remains engaged with R237 (Fig. 5C and 5D). After ADP is released, K184 switches to form an electrostatic interaction with D461 (Fig. 5E) and the lowest energy state m1 in PPS and ADP-bound states is no longer favorable in the rigor state (Fig. 5F). This newly formed interaction between K184 and D461 links ADP release to transducer structural changes. A corresponding salt bridge in myo1b can form upon ADP release as demonstrated in a cryo-EM study (40). In the meantime, the interaction between E179 and R237 becomes weaker (Fig. 5E), as indicated by the metastable state m4 circled by red (Fig. 5F). This destabilization allows the pocket to accommodate an incoming ATP. A previous MD simulation (41) showed that the E179-R237 contact is a unique feature in the nucleotide-favorable state compared with nucleotide-unfavorable states. R237W, a possible DCM-associated mutation (42), has been found to have weaker affinity for nucleotides (11).

## Discussion

By combining comparative modeling and MD simulations, we have investigated the cardiac actomyosin conformational states in the cross-bridge cycle. The approach is unique in the following ways. (i) Comparative modeling by Rosetta enables the initial sampling of conformational ensembles which are difficult to explore via other methods. (ii) Enhanced sampling by GaMD expands the timescale that standard MD can cover. (iii) The modeling and simulations explicitly take the ligand states into account. By comparing the energy profiles of different ligand states, intermediate states and specific residues are found to play important roles during actomyosin force production. (iv) The sequence information can be directly linked to myosin structure and activity. In SI and Fig. S5, we show that our results are more accurate than the rigor actomyosin structures predicted by the AlphaFold-Multimer program (26) (which could not produce PPS structures).

The simulations provide insights into the ordering of events during the force-generation process. By calculating the actin contact area contributions from individual myosin subdomains, we suggested that first the L50 subdomain attaches to actin in the initial binding phase. The HLH motif of L50 could serve as an anchor to facilitate U50 engagement to actin, which is accompanied by the binding cleft closure. This is consistent with previously proposed models for other myosin isoforms, in which L50 binds to actin before U50 (19) and the HLH motif acts as a hinge to allow the rotation of the myosin head (20). Once the L50 subdomain is bound to actin, the cleft closure is coupled to the transition at the motor core, as shown in Fig. 4 by comparing the free energy profile at the actin-bound PPS state with that at the rigor state.

Whether phosphate release precedes the myosin lever arm swing is highly debated in the literature. A crystal structure of MyoVI (39) described an intermediate state (between the PPS and rigor states), in which a “back door” tunnel for Pi release exists between switch II and switch I. These crystals, which were soaked in high levels of phosphate, showed that Pi was able to stick to the exit of the tunnel without changing the structure, or reenter the active site to reverse the myosin back to its PPS state. The results suggested that Pi release acts as a gating event before the major lever arm swing. We note that this structure was solved in the absence of actin and it features an additional relative rotation between L50 and U50 domains compared to both the PPS and rigor states. Our simulations revealed a similar tunnel is formed by switches I and II, even when Pi remains at the active site (Fig. 6C). The salt bridge between R243 (switch I) and E466 (switch II) has been found important in the communication between the actin-binding sites and the active site in mutagenesis studies of other myosin systems (43, 44). It has also been demonstrated that disrupting the switch I – Pi interaction impacts the allosteric communication during the early actin binding phase (45).

Single-molecule optical tweezers results (24, 46) have favored a powerstroke-first model. The estimated time scale of powerstroke for cardiac myosin (1 ms (24)) is independent of Pi concentration and faster than Pi release rate (ranging from a few ms to 60 ms (24, 47)). In these experiments, an external load was applied to myosin and it has been shown that the rate of powerstroke increases by applying a hindering load. A more recent kinetics study proposed a multistep Pi release mechanism in which Pi still leaves the active site before the powerstroke but pauses at a secondary Pi binding site in the tunnel (48). Our results are consistent with this explanation and reveal how different ligand states shift the population of actomyosin conformations. The overlap in the populations of PPS and rigor states shown in the free energy profiles (Fig. 4D and 4E) indicates that actin-bound PPS myosin could transition to a post-powerstroke state without releasing Pi, although with a very small probability. Pi release from the active site shifts the population towards a post-powerstroke state. Adding a load to myosin may impact the rate of Pi and ADP release through adjustments of interactions in the motor core, likely via those allosteric residues involved in Fig. 5 and 6. A follow-up work can be enhanced sampling of the Pi release pathway and studying how the populations of the motor conformations are affected as Pi reaches the surface of the tunnel.

Our approach provides atomistic models that link motor sequence to function. Different myosin isoforms can have distinct cross-bridge kinetics or ordering of events. The structural ensembles obtained here can be used as input templates for efficiently generating distributions for another myosin sequence via a similar protocol. A previous MD simulation (41) with 12 myosin motor domains has demonstrated that the P-loop conformations are well correlated with ADP release rates and duty ratios. Although that work was done for systems of rigor myosin without actin, it highlighted the importance of intrinsic structural ensembles in controlling the mechanochemical cycle. Future applications will focus on predicting how myopathy-related mutations and small molecules impact individual steps in the cross-bridge cycle. For example, an MD simulation (49) showed that the small molecule 2-deoxy-ADP, the hydrolysis product of a myosin activator 2-deoxy-ATP (16, 50), changes the active site conformation compared with the ADP-bound state. Such an effect could be transmitted to the actin-binding region via the key allosteric residues found in this study. Another interesting area would be exploring the effects of other actin-associated proteins such as troponin and tropomyosin.

## Materials and Methods

### Comparative modeling of the myosin-actin complex

The RosettaCM hybridization protocol (33, 51) was used to build structural ensembles based on multiple structure templates (Fig. 1B and 1C). Firstly the query myosin-actin sequence is threaded onto each individual template. Rosetta uses Monte Carlo sampling to produce hybrid-template models by recombing template segments in Cartesian space and de novo building unaligned regions in torsion space. Then the model geometry is further improved by optimizing local structure, e.g. segment boundaries and loops. In this process conformations away from the starting templates are able to be explored through MC sampling with local fragment superposition and energy minimization moves. The high-score models are inspected and selected for MD simulations. Fig. 1C shows the case for the PPS state. For the rigor and ADP-bound states, PDB 5H53 and PDB 6C1D were used as the templates for the myosin-actin complex, respectively. The ATP hydrolysis products (ADP+Pi) were explicitly incorporated at the active site of the PPS myosin. Tropomyosin and troponin were not included in the models. Modeling details are described in SI Appendix.

### Enhanced sampling simulations for the model ensembles

From each ensemble obtained above (PPS, ADP-bound, and rigor states), 35 conformations were selected for the subsequent all-atom simulations. For each Rosetta model, three independent replica runs were launched. Firstly energy minimization and equilibration MD were performed to prepare and filter the model systems for the following GaMD simulations (32). Each GaMD replica consisted of a 10-ns conventional MD stage and a 25-ns GaMD stage. The accumulated GaMD trajectories lasted 2.0 *µ*s, 2.0 *µ*s, and 2.6 *µ*s for the pre-powerstroke, rigor, and ADP-bound states, respectively. For a detailed description of system setup, force field parameters, simulation protocol, and energetic reweighting, see SI Appendix. Representative MD conformations are also provided in Dataset S1.

## ACKNOWLEDGMENTS

This work was supported by National Institutes of Health grant R01-GM031749. The authors gladly acknowledge the computational resources (TSCC (52)) provided by San Diego Supercomputer Center at UCSD. Funding for M.R. was provided by NIH NIAMS P30AR074990 and NIH R01-HL128368.

