## Supporting Information for "Integrating comparative modeling and accelerated simulations reveals conformational and energetic basis of actomyosin force generation"

#### This PDF file includes:

Supporting text
Figs. S1 to S5
Table S1
Legends for Movies S1 to S2
Legend for Dataset S1
SI References

#### Other supporting materials for this manuscript include the following:

Movies S1 to S2
Dataset S1

### Supporting Information Text

### SI Materials and Methods

**Comparative modeling of the myosin-actin complex.** RosettaCM (1) breaks up multiple templates to produce hybridized structures that contain information from different structures. Here we applied the RosettaCM scripts published in Ref (2). A detailed tutorial with examples can be found in the SI of Ref (2). The protocol involves three steps: 1. align the target sequence to templates with Clustal Omega (3) and prepare input files; 2. use the partial thread application to create threaded pdb files for each target-template alignment; 3. generate ensembles of models using the RosettaCM hybridize protocol, which includes three stages of assembly and optimization (1).

The final assembly consisted of one human $\beta$-cardiac myosin (MYH7) and two adjacent $\alpha$-cardiac actin (ACTC1) monomers. For the PPS state modeling, PDBs 5N69 (PPS, bovine cardiac muscle) and 5H53 (rigor, rabbit skeletal muscle) were used as the templates. The cardiac myosin sequence was aligned to the two template sequences and the cardiac actin sequence was aligned to that of 5H53. After partial threading, the threaded pdb of 5N69 included all the atoms of the original pdb; the threaded pdb of 5H53 included the two actin monomers in contact with myosin, as well as the key myosin loop motifs at the interface. For the rigor state modeling, the myosin-actin complex in 5H53 was used as the template. For the ADP-bound modeling, the myosin-actin complex in 6C1D (myosin 1b) was used as the template. The PPS and ADP-bound states have the respective ATP hydrolysis products bound at the active site, which were incorporated by adding the additional flags "-extra_res_fa" and "-extra_res_cen" in the RosettaCM command to load the full-atom and centroid mode ligand parameters. The Rosetta module molfile_to_params.py was used to generate the Pi ($H_2PO_4^-$) and ADP parameters. The ligand states (ADP+Pi for PPS, ADP for ADP-bound, and empty for rigor) were consistent throughout Rosetta modeling and MD simulations. The output models were ranked by the Rosetta score function and 35 models from each ensemble (PPS, ADP-bound, or rigor ensemble) were selected for the following MD simulations.

**MD simulation setup.** All the selected Rosetta models were solvated in a TIP3P water box with 150 mM NaCl. All the MD simulations were performed using the GPU-accelerated version of Amber18 (4, 5) with the ff14SB force field (6). The phosphate ion was modeled as $H_2PO_4^-$, which is the protonation state proposed for the product state of myosin (7, 8). Antechamber and the general AMBER force field (GAFF2) (9, 10) were applied to assign bonded and LJ parameters for Pi, whose partial charges were assigned according to Ref (11). An existing set of ADP parameters (12) and a multisite $Mg^{2+}$ model (13) were used. Amber's tleap program was employed to generate the input files.

**Simulation protocol.** For each MD system built on a Rosetta model, three independent replica runs were performed, as described below. Firstly energy minimization was carried out while constraining the protein atom positions using harmonic potentials with a spring constant of 1 kcal/(mol Å$^2$). With the same positional constraints, a following 10 ns equilibration simulation was performed at 300 K. To maintain the temperature, Langevin dynamics with a friction coefficient of 1 ps$^{-1}$ was applied. Particle Mesh Ewald (14) was used for full-system periodic electrostatics while a 9 Å cutoff was applied to Lennard-Jones interactions. Bonds involving hydrogen atoms were constrained using the SHAKE algorithm (15). The initial simulations filtered out a few systems that resulted in unstable dynamics. We then ran a subsequent GaMD simulation (16) for each system that passed the initial equilibration stage.

GaMD accelerates the sampling of protein conformational transitions by reducing the energy barrier with a harmonic boost potential (17). It has been successful in studying a few molecular machines, such as GPCR (18), CRISPR-Cas9 (19), and $\gamma$-secretase (20). Here the GaMD module implemented in Amber (16) was employed to perform the simulations, which included a 10-ns conventional MD stage and a 25-ns GaMD stage in the isothermal-isobaric ensemble at 1 bar and 300 K. The 10-ns conventional MD was used to collect statistics for calculating GaMD acceleration parameters. In the GaMD stage, the total potential energy surface was smoothed by a boost potential that had a 6 kcal/mol upper limit of the standard deviation for accurate reweighting. MC barostat (21) was chosen for pressure control. The accumulated GaMD trajectories lasted 2.0 $\mu$s, 2.0 $\mu$s, and 2.6 $\mu$s for the pre-powerstroke, rigor, and ADP-bound states, respectively.

**Data analyses.** To obtain two-dimensional (2D) free energy profiles from the GaMD runs, we construct a 2D histogram with a total number of $M$ bins, which cover the 2D space of interest. We define $\delta_{m,i}$ as the indicator function (22) for the $i$th frame of the trajectory through

$$\delta_{m,i} = \begin{cases} = 1 & \text{if frame } i \text{ falls in bin } m \\ = 0 & \text{otherwise} \end{cases} \qquad [1]$$

The weighted histogram at bin $m$ can be computed by

$$H_m = \sum_{i=1}^{N} \delta_{m,i} e^{\beta \Delta V_i} \qquad [2]$$

where $\Delta V_i$ is the boost potential at the $i$th frame, $N$ is the total number of frames, and $\beta = (k_B T)^{-1}$. The Maclaurin series expansion method was used to approximate the exponential term $e^{\beta \Delta V_i}$ (22). One can then determine the 2D free energy profile via

$$F_m = -k_B T \log H_m + F_0 \qquad [3]$$

where $F_0$ is an arbitrary constant which is chosen here to set the minimum value in the free energy profile to zero. 1D free energy profiles can be obtained following a similar approach.

Movies 1 and 2 were rendered with VMD (23).

### Additional analyses on the actin-myosin interactions at the pre-powerstroke state.

Due to the weak binding nature of the PPS state, experimentally it is difficult to characterize this state. We have shown that the PPS state is represented by an ensemble of conformations rather than a single conformation (Fig. S3). For the PPS simulations, we were able to find a small portion of frames in which the CM loop and loop 4 are disengaged from actin. There could exist some very weak binding states during which the myosin diffuses on the actin surface to search for the specific binding site, but these myosin-actin interactions are likely non-specific and not included in this study. Compared to the rigor state (Fig. S3A and S3C), the PPS state (Fig. S3B and S3D) has a much more versatile interface. For example, two metastable states highlighted in red circles in Fig. S3B show very small contact areas between the cardiomyopathy loop and actin, although the HLH motif remains attached to actin.

Fig. S4 demonstrates the distributions of the distances between K635 (myosin) and two negatively charged residues E4 and E3 (actin) in the rigor (Fig. S4A) and PPS (Fig. S4B) simulations. In either the rigor or PPS simulations, the most probable state is when K635 forms salt bridges with E4 or E3 on actin (distance$\sim$3.8Å); however, the second peak becomes relatively more populated at the PPS state than the rigor state. For example, for the distance K635-E3 (cyan curves), the relative probability density between the first and second peaks is 1:0.63 for the rigor state, whereas for the PPS state, this ratio becomes 1:0.94. Consistent with Fig. S3D, this observation shows that the interactions between the actin N-terminus and loop 2 are more versatile at the PPS state.

### AlphaFold2 modeling of the actomyosin complex

To see if AlphaFold can predict the structure for the actin-myosin complex, we applied the AlphaFold-Multimer module (24) in the AlphaFold2 package (25) (https://github.com/deepmind/alphafold). The following settings were used: model_preset=multimer, db_preset=reduced_dbs, max_template_date=2020-01-01, mgnify_database_path=mgy_clusters_2018_12.fa, use_gpu_relax=true. For a given input, 5 seeds were generated for each of the five AlphaFold2 models, resulting in 25 total predictions. The LDDT (pLDDT) scores (25) were used to rank model confidence.

Here, two different input systems were tested: one system (1myosin:2actin) contains one myosin monomer and two actin monomers; the other one (1myosin:1actin) contains one myosin monomer and one actin monomer. Interestingly, the input with 1myosin:2actin could not produce a rigor interface, whereas the input with 1myosin:1actin only produced three “rigor-like” complexes out of the 25 models. The top five models for the 1myosin:1actin system are demonstrated in Fig. S5A (PDBs also provided in Dataset S1). All the myosin molecules are in the rigor state. By measuring the distance between the CB atoms of V406 (CM loop) and A26 (actin), we find only Models 2 and 4 represent tight interfaces with such a distance smaller than 5.5 Å. The actin binds to the wrong side of myosin in Models 1 and 3; the CM loop and HLH motif are away from the correct docking site in Model 5. Model 2 has the closest conformation to a rigor actomyosin structure among all top models, but its lower 50k domain (such as the HLH motif) is not docked to the correct position on actin (Fig. S5B). So our protocol generates better rigor models than the ones predicted by AlphaFold2. We note that AlphaFold2 was unable to generate a pre-powerstroke model.

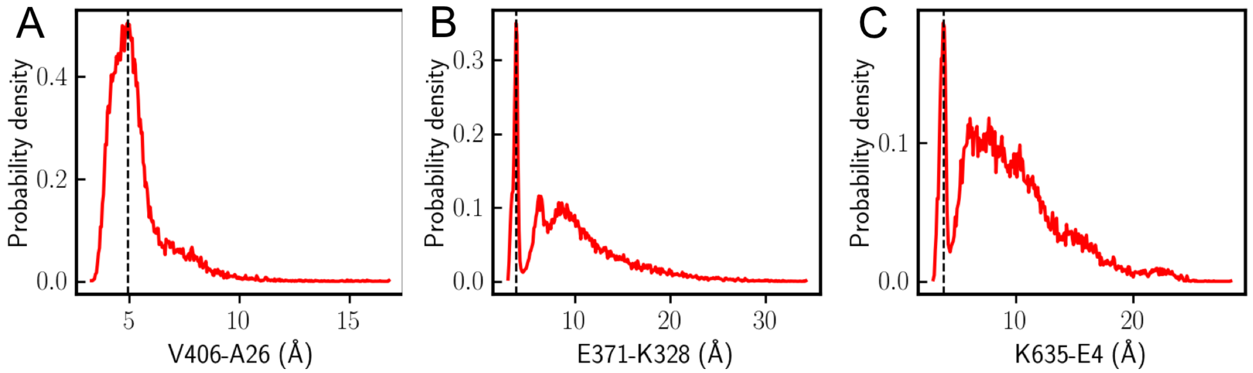


**Fig. S1.** Probability distributions of distances between key residue pairs at the myosin-actin interface in the rigor state: A. the distance between the V406 and A26 (CB atoms); B. the distance between E371 (CD atom) and K328 (NZ atom); C. the distance between K635 (NZ atom) and E4 (CD atom). The dashed black lines show the most probable distances.

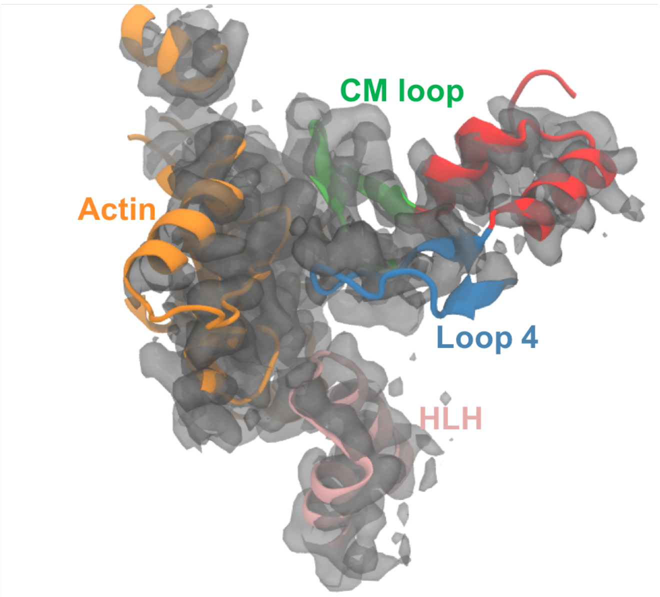


**Fig. S2.** Fitting key myosin-actin interface motifs into a cryo-EM density of the rigor cardiac actomyosin complex (EMD-22335, 3.8 Å resolution).

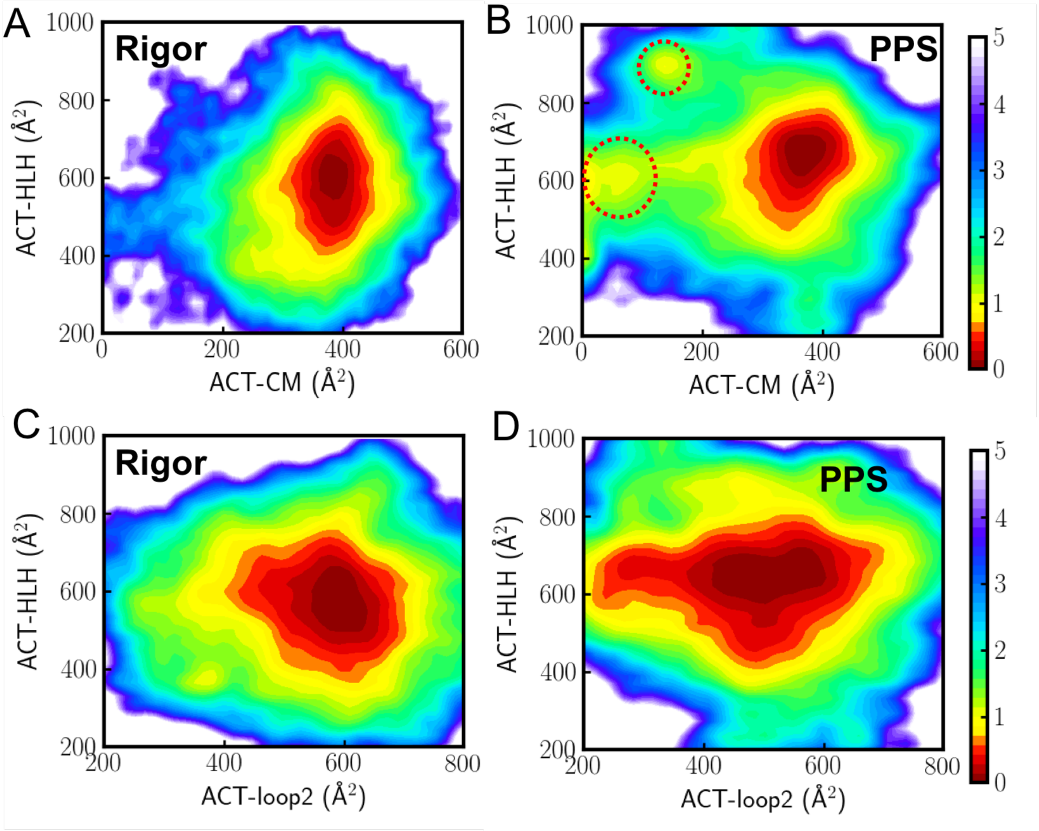


**Fig. S3.** 2D free energy profiles projected onto two contact area coordinates in the rigor (A and C) and PPS states (B and D). In panels A (rigor) and B (PPS), the x-axis represents the contact area between actin and the CM loop, whereas the y-axis represents the contact area between actin and the HLH motif. The red circles in B highlight two metastable states, in which the CM loop has relatively smaller contact areas with actin, while the HLH motif remains closely engaged with actin. In panels C (rigor) and D (PPS), the x-axis represents the contact area between actin and loop 2; the y-axis represents the area between actin and the HLH motif.

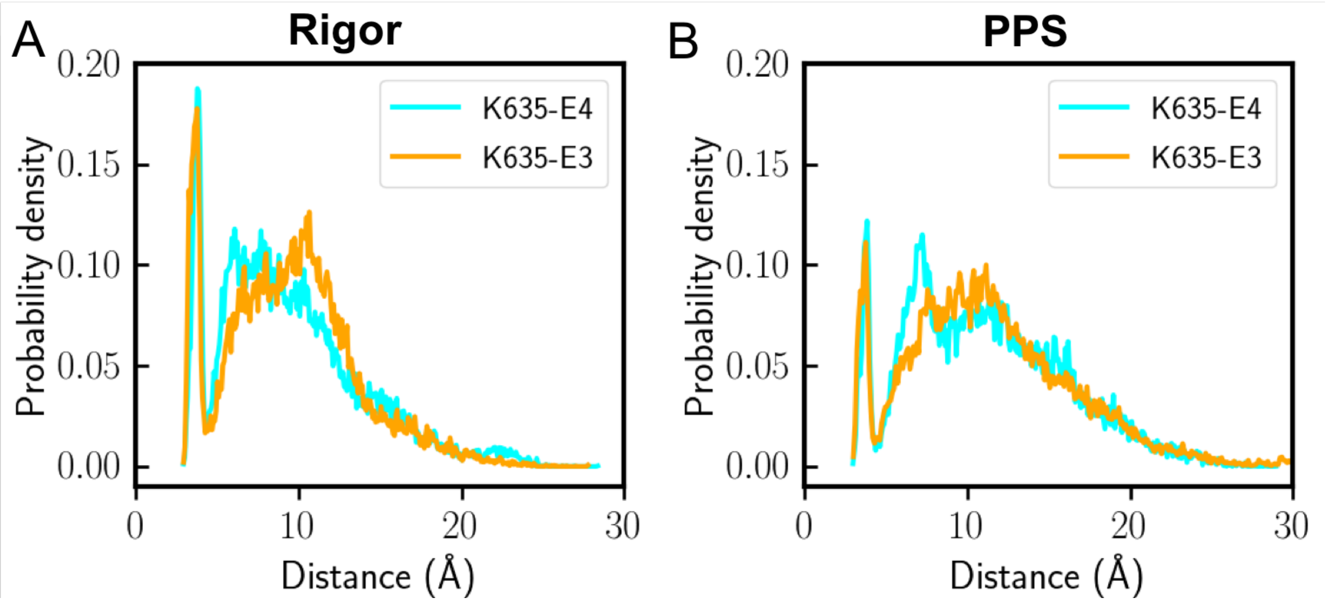


**Fig. S4.** Measuring the population distributions of key distances between actin and myosin loop 2. A. The distributions of the distances between myosin K635 and actin E4/E3 are measured for the rigor simulations. B. Same distance distributions are measured for the PPS simulations.

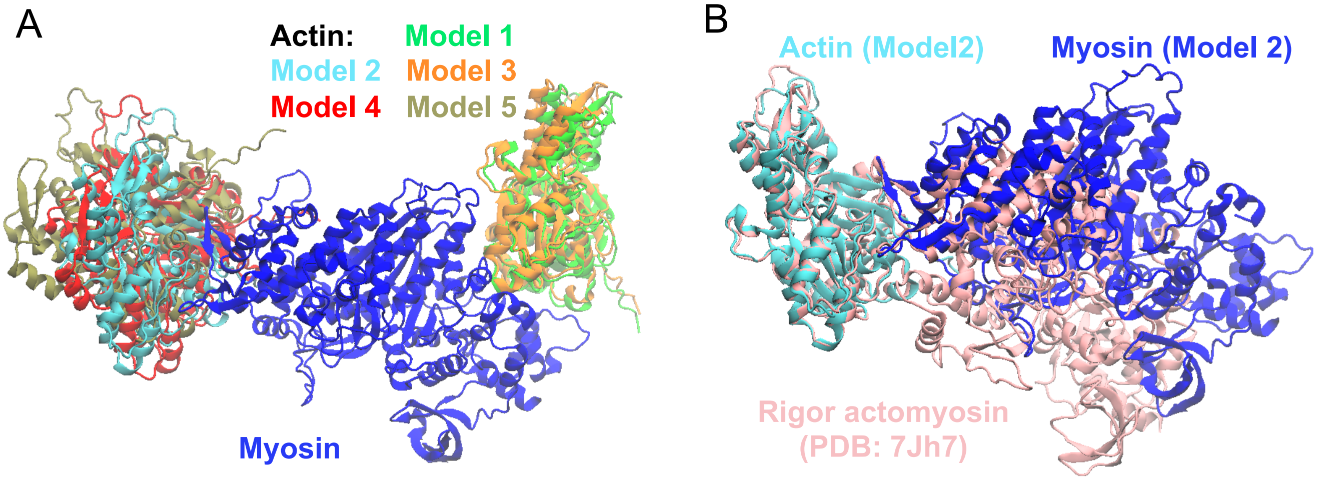


**Fig. S5.** A. Top five AlphaFold models (Models 1-5) for the myosin-actin complex (1myosin:1actin). The complex structures are aligned by myosin. Only Models 2 and 4 resemble tight interfaces somewhat similar to the rigor interface. Since the individual conformations of myosin (rigor) are very similar among the top 5 models, only the myosin model of Model 2 is displayed (blue). B. Model 2 is aligned to the experimental structure of the rigor cardiac actomyosin complex (PDB 7JH7, colored in pink).

**Table S1. Averaged MD RMSD[a] values from experimental PDBs at different myosin states**

| Simulation system | 7JH7 (26) Rigor actomyosin | 6X5Z (27) Rigor actomyosin | 5N69 (28) PPS myosin |
|---|---|---|---|
| Rigor | 2.30±0.29 Å | 2.31±0.28 Å | 3.58±0.30 Å |
| PPS | 3.50±0.22 Å | 3.45±0.23 Å | 1.52±0.25 Å |

a. RMSD is calculated using the $C_\alpha$ atoms of the motor domain (residues 95 to 705, excluding the flexible loops on the surface). Three PDBs are used as the reference structures for RMSD calculations: 7JH7, 6X5Z, and 5N69.

**Movie S1. The actomyosin conformational dynamics at the rigor state. Loop 2, CM loop, loop 4, myosin motor domain, and actin are colored blue, lime, cyan, red and gray, respectively.**

**Movie S2. A gate is formed between switch I (yellow) and switch II (purple). P loop is colored green. The gating residues (R243 and E466) and Pi/ADP/$Mg^{2+}$ are shown in atomic representation. Another metastable salt bridge (E525-K484) is shown in the lower part of the movie.**

**SI Dataset S1 (Rep_Conformations.zip)**

This file includes two top representative conformations for both the rigor and PPS simulations (after clustering analysis). Top five AlphaFold models are also provided.
